# Oncogenic role of sFRP2 in P53-mutant osteosarcoma development via autocrine and paracrine mechanism

**DOI:** 10.1101/246454

**Authors:** Huen Suk Kim, Seungyeul Yoo, Jeffrey M. Bernitz, Ye Yuan, Andreia M. Gomes, Michael G. Daniel, Jie Su, Elizabeth G. Demicco, Jun Zhu, Kateri A. Moore, Dung-Fang Lee, Ihor R. Lemischka, Christoph Schaniel

## Abstract

Osteosarcoma (OS), the most common primary bone tumor, is highly metastatic with high chemotherapeutic resistance and poor survival rates. Using induced pluripotent stem cells (iPSCs) generated from Li-Fraumeni syndrome (LFS) patients, we investigated an oncogenic role of secreted frizzled-related protein 2 (sFRP2) in P53 mutation-associated OS development. Interestingly, we found that high sFRP2 expression in OS patient samples correlates with poor survival. Systems-level analyses identified that expression of sFRP2 increases during LFS OS development and can induce angiogenesis. Ectopic sFRP2 overexpression in normal osteoblast precursors is sufficient to suppress normal osteoblast differentiation and to promote OS phenotypes through induction of oncogenic molecules such as FOXM1 and CYR61 in a β-catenin independent manner. Conversely, inhibition of sFRP2, FOXM1 or CYR61 represses the tumorigenic potential. In summary, these findings demonstrate the oncogenic role of sFRP2 in P53 mutation-associated OS development and that inhibition of sFRP2 is a potential therapeutic strategy.

## Introduction

Osteosarcoma (OS) is the most common primary bone tumor. It accounts for about 60% of all primary bone tumors and about 2% of all childhood cancers (Kansara et al, 2014; Reed et al, 2017). Despite significant advancements in OS treatment modalities, including surgery, chemotherapy and targeted drug therapies, the five-year overall survival rate has remained stable over the last 20 years at about 65% for patients with primary OS and less than 20% for patients with metastasis (Bielack et al, 2016; Jaffe et al, 2013). This stagnation of clinical outcomes underlines the urgent necessity of novel model systems to study the mechanism of OS development in a patient-specific context and to identify molecular targets for the development of new therapeutic strategies.

P53, the most studied tumor suppressor, regulates cell cycle, apoptosis, senescence, metabolism, and cell differentiation (Bieging et al, 2014; Laptenko & Prives, 2006; Lee et al, 2012; Vogelstein et al, 2000). Therefore, it is no surprise that aberrant P53 expression contributes significantly to cancer development (Brosh & Rotter, 2009; Goh et al, 2011; Muller & Vousden, 2014; Petitjean et al, 2007). Half of all human sporadic bone tumors have genetic lesions in *TP53* (Hung & Anderson, 1997; Reed & Shokat, 2014; Toguchida et al, 1992). Patients with Li-Fraumeni syndrome (LFS), which is caused by mutations in *TP53*, show a 500-fold higher incidence of OS relative to the general population (Birch et al, 1994; Hisada et al, 1998). Genetic manipulation of p53 function in mice confirmed the significance of P53 in OS development (Iwakuma & Lozano, 2007; Jacks et al, 1994; Lozano, 2010) and identified mesenchymal stem cells (MSCs) and pre-osteoblasts (pre-OBs) as the cellular origin of OS (Rubio et al, 2014; Velletri et al, 2016). OB-restricted deletion of p53/MDM2 or p53/Rb in mice resulted in OS development at a high penetrance of about 60% and 100%, respectively (Lengner et al, 2006; Walkley et al, 2008).

Secreted frizzled-related protein 2 (sFRP2), widely known as a WNT antagonist, was initially reported as a secreted anti-apoptosis related protein (Melkonyan et al, 1997; Polakis, 2000). Indeed, ectopic expression of sFRP2 promotes cell growth and has antiapoptotic properties in renal and breast cancer (Lee et al, 2004a; Lee et al, 2004b; Yamamura et al, 2010). Conversely, sFRP2 hypermethylation and its decreased expression have been associated with prostate, liver and gastric cancer (Cheng et al, 2007; O'Hurley et al, 2011; Perry et al, 2013; Takagi et al, 2008). The role of sFRP2 in cancer is still controversial.

Using LFS patient-derived induced pluripotent stem cells (iPSCs), we previously recapitulated the pathophysiological features of LFS-mediated OS development in vitro and in vivo (Lee et al, 2015). Taking advantage of this platform, we observed increased expression of sFRP2 during LFS iPSC-derived OB differentiation. As a result of these findings and because the exact function of sFPR2 in OS is not clear, we investigated its role in LFS-mediated abnormal OB differentiation, tumorigenesis, and OS development. Here, we report that sFRP2 overexpression (sFRP2 OE) induces OS phenotypes increasing FOXM1 expression, promotes angiogenesis and endothelial expression of the matricellular protein CYR61. Conversely, targeting sFRP2 OE in LFS and OS has therapeutic promise for subtypes of OS with P53 mutation.

## Results

### sFRP2 is overexpressed in LFS (G245D) iPSC-derived OBs

In order to discover potential therapeutic targets for LFS (G245D) mediated OS, we analyzed the genome-wide transcripts of our LFS data set (GSE58123) composed of LFS iPSC-derived MSCs differentiated to OBs, specifically MSCs (D0) and pre-OBs at D7, D14 and D17 of *in vitro* differentiation (Lee et al, 2015), and compared it with OS GEO data sets (GSE33458, OS tumor-initiating cells) (Fig 1A) (Rainusso et al, 2011). Differentially expressed genes (DEGs) were sorted with the time-independent t-test (p<0.05). This method enables extraction of the most significantly and consistently up- or downregulated genes during *in vitro* OB differentiation in the LFS dataset (Fig 1A). We identified sFRP2 as an overexpressed gene with potential implication in OS development. Quantitative PCR and western blot analyses confirmed that sFRP2 expression is significantly overexpressed in LFS iPSC-derived OBs compared to control OBs (Fig 1B). The information of LFS iPSC-derived OBs that we used in this study are summarized in Supplementary Table S1. sFRP2 expression was also detected in several OS cell lines harboring p53 mutations (HOS, HOS/MNNG, 143B) or MDM2 amplification (SJSA-1) (Fig 1C), and in subcutaneous tumors obtained from mouse xenograft assays with LFS cell lines (Lee et al, 2015)(Fig 1D). In order to confirm the correlation between the *TP53* mutation and sFRP2 expression in MSCs, we introduced the P53 (G245D) mutation into WT MSCs using CRISPR-Cas9 (Supplementary Fig S1A). We observed that sFRP2 expression was significantly increased in WT MSCs edited to harbor the P53 (G245D) mutation, but not in WT MSCs with P53 depletion (shRNAs to P53) (Supplementary Fig S1B). Fibroblasts from LFS patients with a G245D mutation also had higher sFRP2 expression (Supplementary Fig S1C). These results demonstrate that sFRP2 OE is induced by P53 mutation (G245D).

**Figure 1.**
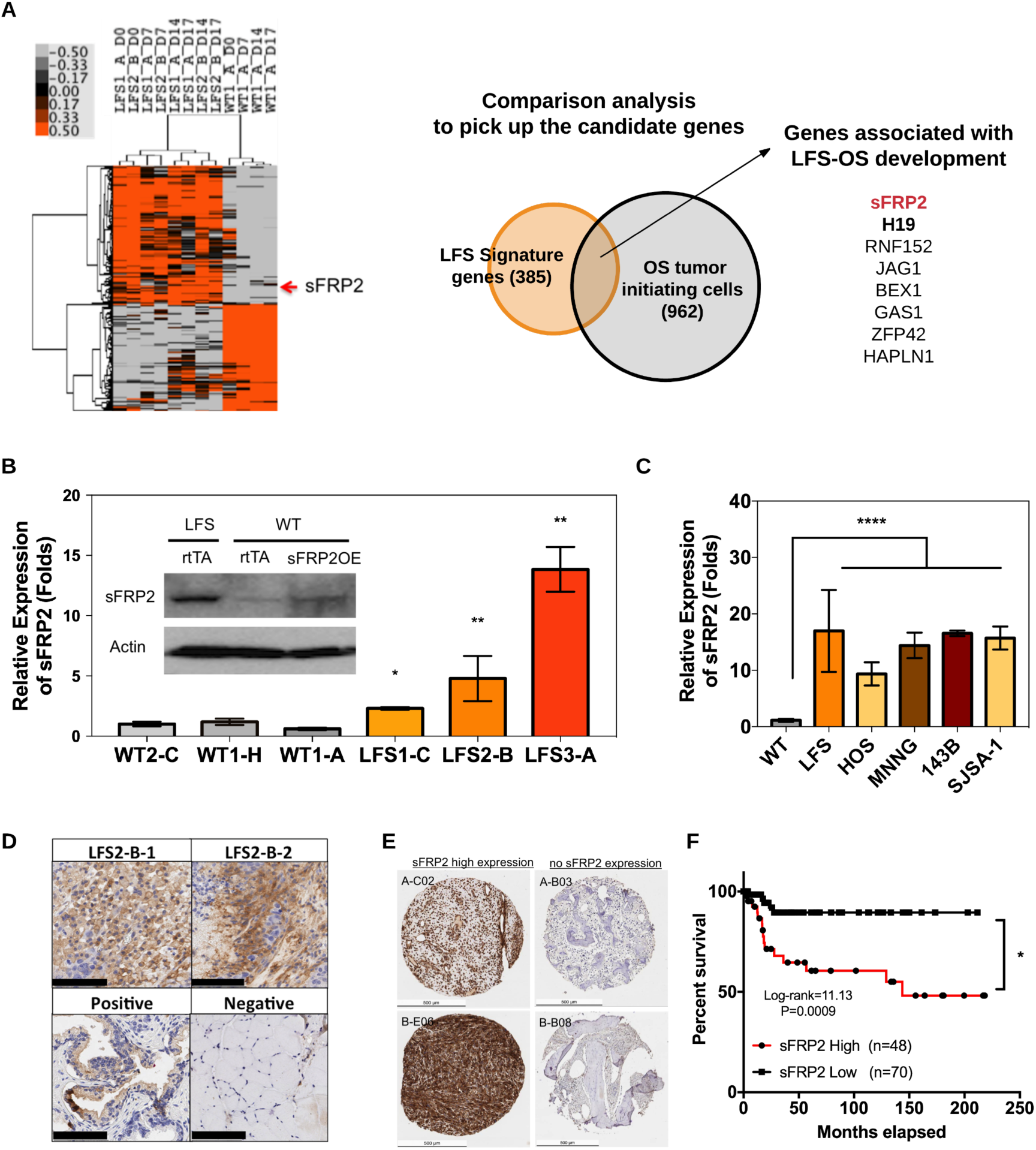
sFRP2 OE in LFS (G245D) mediated OS. **A.** Global transcriptome analysis of LFS and WT iPSCs-derived MCS differentiated to OBs. Differentially expressed genes (DEGs) were sorted by time-independent t-test. sFRP2 is an overexpressed gene that is also enriched in the signature gene list of an OS gene set (GSE33458). **B.** Quantitative PCR analysis and western blotting of sFRP2 expression (D4 of differentiation; mean±SD; n=3 independent repeats in triplicates) in LFS and WT MSCs (*, p-value< 0.05; **, p-value < 0.0001). **C.** sFRP2 expression in human OS cell lines. (Data are shown as n=3 independent repeats in triplicates; ****, p-value< 0.0001, ANOVA). **D.** Expression of sFRP2 protein in LFS subcutaneous xenograft tumors. Human prostate tissue was used as positive and mouse muscle tissue as negative staining control (Scale bar = 100 µm). **E.** Representative staining of sFRP2 protein expression levels in OS tissue samples. The tissue microarray contained 151 OS patient samples. **F.** Survival analysis of OS patients linked to the tissue microarray. Survival curve of sFRP2 high and low expression groups was calculated using the log-rank test (chi-square= 11.13, p-value=0.0009).

Next, we analyzed the expression of sFRP2 levels in human OS tissue samples using a tissue microarray spotted with 151 OS patient tissue samples (Figs 1E, and S1D). Unfortunately, there is no data on P53 status associated with this OS tissue array. Nevertheless, high sFRP2 expression (intensity score 2 and 3, with 3 being the highest expression) negatively correlated with patient survival (Figs 1F and Supplementary S1E, and Supplementary Table S2). There was no gender difference in tumor incidence (Supplementary Fig S1F) and the area staining positive for sFRP2 is small in the group with low sFRP2 expression (Supplementary Fig S1G). These data indicate that sFPR2 OE is linked to the P53 mutation in LFS OS patients and associated with poor prognosis for sporadic OS patients.

### Global gene expression analysis of sFRP2 OE

In order to better understand the molecular role of sFRP2 OE in LFS cells, we generated global gene expression profiles of WT cells overexpressing sFRP2 (sFRP2OE), LFS cells with sFRP2 knock down (sFRP2KD), parental WT (WT2-C) and LFS cells (LFS2-B) at four time points during *in vitro* OB differentiation (D0, D7, D14 and D17) (Supplementary Fig S2A). We used stable cell lines with sFRP2 OE or KD (Supplementary Fig S2B). Principal component analysis (PCA) showed that WT and LFS cells are clearly distinct from each other (Fig 2A), consistent with our previous LFS RNA-seq data (Lee et al, 2015). The expression patterns of sFRP2OE were also distinct from WT and more evident at later stages of differentiation (Fig 2A). Likewise, LFS data sets diverged more strongly at late stages of differentiation, while sFRP2KD data clustered together and were distinct from LFS groups (Fig 2A). This result shows that sFRP2 OE significantly alters gene expression in pre-OBs during differentiation. Linear regression with conditioning time points identified DEGs between sFRP2OE and WT, between LFS and WT, and between sFRP2KD and LFS groups (p<0.001, corresponding permutation FDR<0.01) (Fig 2B). DAVID functional annotation was used to understand the general biological effect of DEGs associated with sFRP2 OE. WT signature genes (DEGs during OB differentiation) were generally involved in bone mineralization and BMP and WNT signaling pathways required for normal OB differentiation (Fig 2C). Interestingly, sFRP2OE signature genes were linked to cell proliferation, cell adhesion, and cell migration, categories implicated in the tumorigenic role of sFRP2 (Kaur et al, 2016; Lee et al, 2004b; Siamakpour-Reihani et al, 2011) whereas, sFRP2KD signature genes restored bone mineralization, apoptosis process, and negative regulation of cell proliferation (Fig 2C). Mouse Gene Atlas analysis of DEGs from WT, sFRP2OE and sFRP2KD signatures confirmed that sFRP2 OE significantly affects differentiation, while sFRP2 KD recovered normal OB differentiation expression signatures (Supplementary Fig S2C).

**Figure 2.**
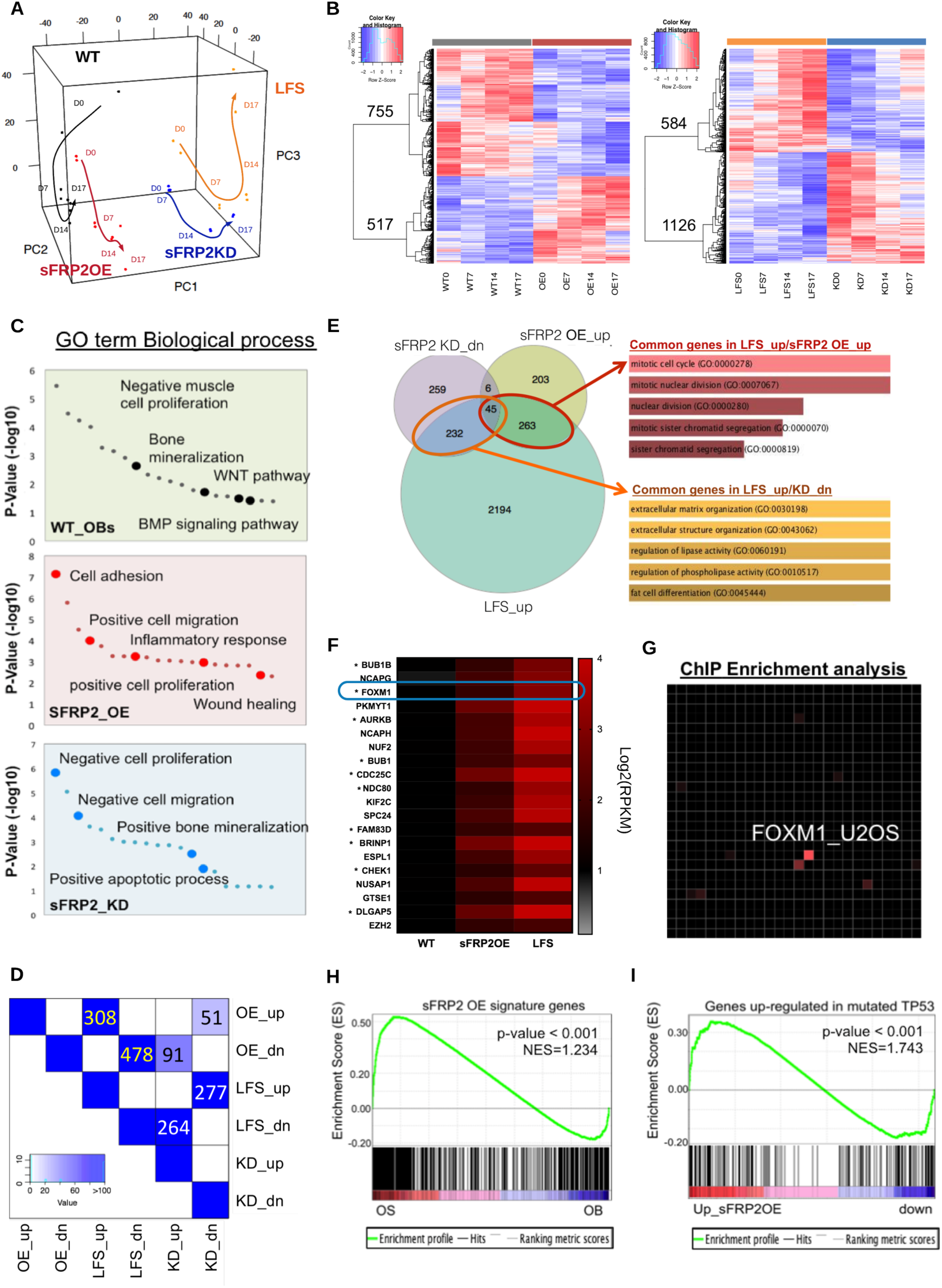
Global gene expression analysis reveals that sFRP2 OE induces cell proliferation and perturbs the differentiation process. **A.** PCA of RNA-seq data from sFRP2 OE and KD samples (variance; PC1=0.4020, PC2=0.1839, PC3=0.0903). **B.** Heatmaps of DEGs in sFRP2 OE and KD samples (DEG cutoff p-value<0.001). **C.** DAVID GO term biological process analysis of DEGs from WT, sFRP2 OE, and sFRP2 KD. **D.** Overlaps of DEGs among sFRP2 OE, KD, and LFS cells. Numbers in blue boxes indicate the number of overlapping genes. **E.** Enrichr GO biological process analysis of common genes among sFRP2 OE upregulated genes, LFS upregulated genes, and KD downregulated genes. **F.** Relative expression of cell cycle and cell proliferation-related DEGs commonly upregulated in sFRP2OE and LFS cells (Asterisks, cancer-associated genes). **G.** Common upregulated DEGs are enriched in FOXM1_U2OS ChIP data (Enrichr). (H) GSEA of sFRP2 OE signature with OS gene set (GSE36001). (I) GSEA of sFRP2 OE signature with *TP53* mutation gene set (MSigDB C6 analysis).

In order to determine whether sFRP2OE is significantly associated with oncogenic molecular features of LFS, we compared sFRP2OE to LFS and sFRP2KD signature genes. There were 308 upregulated and 478 downregulated genes common between sFRP2OE and LFS expression signatures (Fisher’s Exact Test (FET) p-value = 1.4×10^−171^ and 2.27×10^−160^, respectively), while 277 downregulated and 264 upregulated genes were reversely overlapped between sFRP2KD and LFS expression signatures (FET p-value of 2.13×10^−130^ and 2.8×10^−127^, respectively) (Fig 2D and Supplementary Table S3). The most common upregulated genes between sFRP2OE and LFS groups include cell cycle and well-known oncogenic genes frequently found in cancers such as AURKB, BUB1B, and FOXM1 (Figs 2E and F). Interestingly, Enrichr (Chen et al, 2013; Kuleshov et al, 2016) ChIP enrichment analysis (Lachmann et al, 2010) revealed that these common upregulated genes in LFS/sFRP2OE were enriched for FOXM1 targets based on prior ChIP-seq data collected from the human OS cell line U2OS (p-value = 1.52×10^−16^) (Fig 2G). This suggests correlation between sFRP2 OE-mediated cell proliferation and oncogenic function of FOXM1 target genes in human OS. Common downregulated genes in LFS/sFRP2OE are associated with extracellular matrix and cell-cell adhesions (Supplementary Fig S2D). Interestingly, tumor suppressor genes such as CADM1 and CADM4 (Tumor suppressor gene database (Zhao et al, 2016), FET p-value=0.004) were enriched in the downregulated gene set (Supplementary Fig S2D).

Finally, we compared sFRP2OE DEGs to the human OS GEO set (GSE36001) in order to determine the correlation with OS features through gene set enrichment analysis (GSEA). sFRP2OE and LFS DEGs were highly correlated with human OS signature genes, whereas sFRP2KD and WT DEGs were highly concordant with OB signature genes (Figs 2H and Supplementary S2E). Additionally, sFRP2OE signature genes were significantly enriched in upregulated genes of the NCI-60 panel of cell lines harboring *TP53* mutations and that of the *Rb1* knockout mouse, suggesting that sFRP2 strongly contributes to the signature of well-known OS genetic lesions, RB1 deletion and P53 mutation (Figs 2I and Supplementary S2F). Collectively, our global transcriptome analyses revealed osteosarcomagenic properties of sFRP2 OE in LFS and OS.

### sFRP2 OE dysregulates OB differentiation through BMP/WNT suppression

We previously reported that LFS iPSC-derived MSCs and pre-OBs have defects in OB differentiation, supporting the idea that compromised P53 dysregulates OB differentiation and promotes generation of abnormal MSCs and pre-OBs, the cells of origin of OS (Lee et al, 2015). Because our global transcriptome analyses provided evidence that sFRP2 OE potentially perturbs OB differentiation (Fig 2C), we wondered if sFRP2 OE in LFS suppresses normal OB differentiation as an oncogenic downstream modulator of mutant P53. We performed alkaline phosphatase (AP) and Alizarin staining on LFS, sFRP2OE, and WT pre-OBs at different differentiation time points (D0-D21). Normal OBs generally exert a high level of AP activity and secrete large amounts of calcium (mineral deposition) detected by Alizarin staining (Savkovic et al, 2014; Yang et al, 2013; Yeo et al, 2007). sFRP2 OE in WT cells clearly suppressed normal OB differentiation similar to LFS cells (Figs 3A and B). In contrast, sFRP2 KD rescued osteogenic AP activity and mineral deposition (Supplementary Figs S3A and B). In agreement with this observation, LFS and sFRP2OE OBs had lower levels of pre-OB markers, (AP/ALPL and COL1A1), mature OB markers (BGLAP/Osteocalcin and PTH1R), osteocyte markers (ZNF521 and FGF23) and OB differentiation transcriptional factors (RUNX2 and Osterix) (Fig 3C). Interestingly, ATF4, whose expression is high in OS tissue and OS cell lines (Zeng et al, 2016), was upregulated in sFRP2OE and LFS (Fig 3C). These results indicate that sFRP2 OE as a consequence of mutant P53 perturbs normal osteogenic processes, which is consistent with the RNA-seq biological process analysis (Fig 2C).

**Figure 3.**
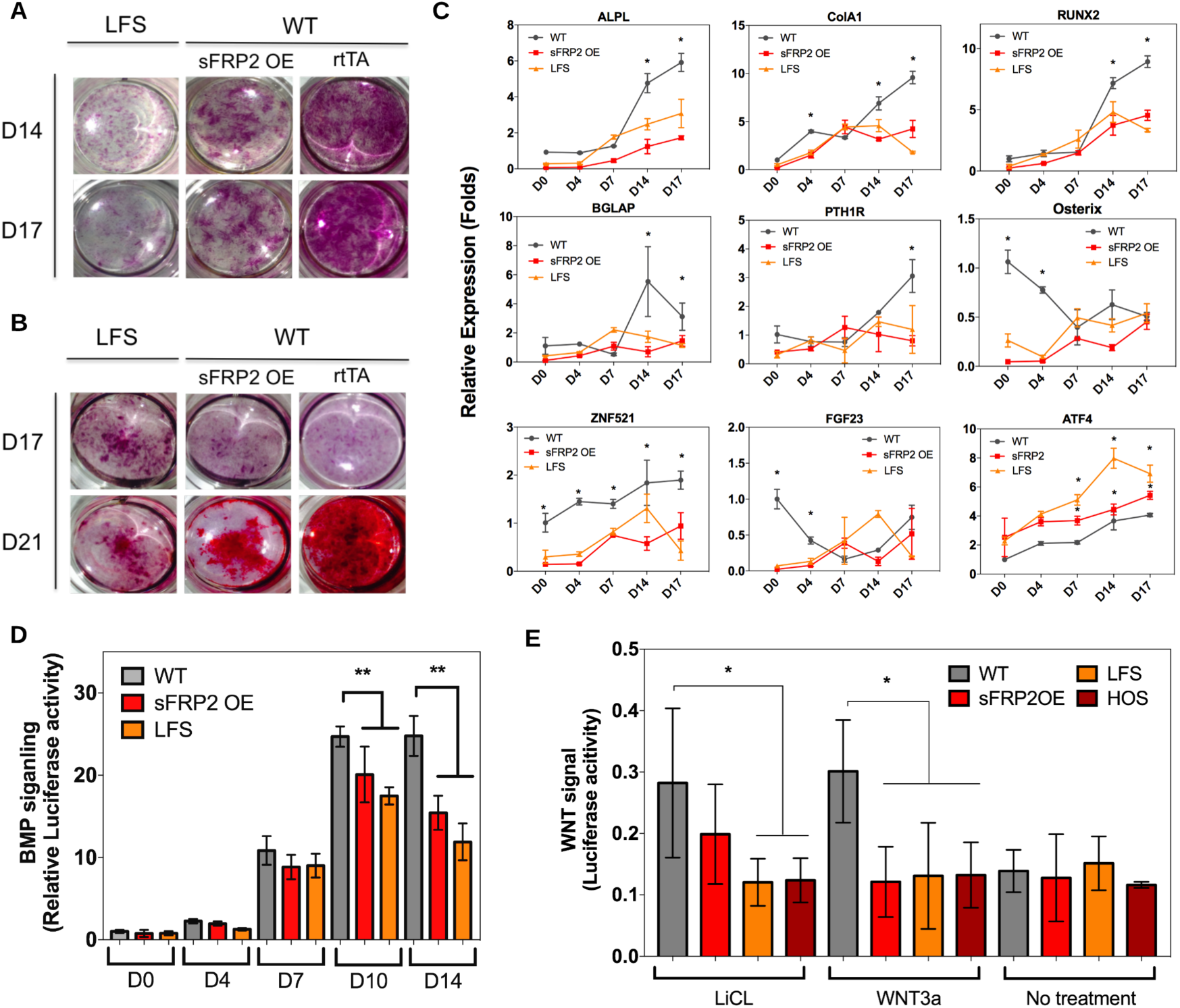
sFRP2 OE suppresses OB differentiation in vitro. **A.** Osteogenic AP analysis in LFS, sFRP2OE and WT cells. **B.** Alizarin staining in LFS, sFRP2OE, and WT derived pre-osteoblast cells. **C.** Quantitative PCR analysis of indicated osteogenic markers (mean±SD, n=3 independent repeats in triplicates, ANOVA) in LFS, sFRP2OE and WT cells (*, p-value< 0.05). **D.** BMP signaling activity in LFS, sFRP2OE and WT cells as measured by a BMP reporter (pSBE3-Luc) **E.** WNT signaling activity as measured by TOP/FOP flash luciferase reporter. Activity levels were normalized using an internal control (dual luciferase assay system). Data are represented as mean±SD, n=3 independent repeats in triplicates (* p-value < 0.05, **, p-value< 0.0001, ANOVA analysis).

The BMP/SMAD and canonical WNT pathways are required for normal osteogenesis (Long, 2011; Noel et al, 2004; Rahman et al, 2015; Song et al, 2012). Particularly, BMP ligands facilitate OB differentiation through the BMP/SMAD pathway (Bandyopadhyay et al, 2006; Kamiya et al, 2008; Noel et al, 2004; Rahman et al, 2015). Using a luciferase-based phospho-SMAD reporter system we monitored BMP signaling activity throughout *in vitro* differentiation (D0-D14). While BMP signaling was relatively low in MSCs and progenitors (Day 4), it increased in pre-OBs (D10 to D14) (Fig 3D). This result is consistent with published *in vivo* data (Abzhanov et al, 2007; Bandyopadhyay et al, 2006). As expected, sFRP2OE and LFS cells showed lower BMP signaling activity than WT OBs. It supports the idea that sFRP2 OE perturbs normal OB differentiation in part through inhibition of the BMP pathway.

The effects of sFRP2 on canonical WNT signaling are context dependent. Generally, sFRP2 suppresses WNT signaling by binding WNT ligands, thus preventing their binding to WNT receptors (Alfaro et al, 2010; Roth et al, 2000; Suzuki et al, 2004). However, sFRP2 has also been shown to activate WNT signaling at low concentrations or in certain cell types (Lin et al, 2016; Sugiyama et al, 2013; Xavier et al, 2014; Yamamura et al, 2010). To understand the role of sFRP2 in canonical WNT signaling in OS development, we performed western blot analysis and β-catenin staining. Compared to WT cells, LFS, sFRP2OE, and HOS cells consistently had lower levels of cytoplasmic β-catenin (Supplementary Fig S3C) while the amount of nuclear β-catenin only increased in WT groups in the WNT3a treatment (Supplementary Fig S3D). Using the TOP/FOP flash luciferase WNT signaling reporter system, we observed low basal WNT activity in all groups (Fig 3E, no treatment). We next activated WNT signaling using WNT3a (100ng/ml) or LiCl (10mM), which inhibits GSKβ. sFRP2OE showed a reduction in WNT3a mediated canonical WNT activity (Fig 3E). LFS and OS cells (HOS) exhibited little or no response to either LiCl or WNT3a (Figs 3E and Supplementary S3E). Ectopic expression of constitutively active β-catenin (DeltaN90) in LFS did not facilitate cell proliferation nor promote cell transformation (Fig. S3 F). However, KD of β-catenin led to severe cell death of LFS cells, confirming that β-catenin is necessary for homeostasis and viability of MSCs and OBs (Supplementary Fig S3G)(Georgiou et al, 2012; Hay et al, 2014). Although it is widely accepted that hyper WNT signaling is associated with oncogenicity in many cancers, our data support that hyper WNT is not involved in osteosarcomagenesis. This is consistent with the observation of inactive canonical WNT activity in high-grade human OS (Cai et al, 2010).

### sFRP2 OE transforms pre-OBs into OS-like cells

We next investigated the oncogenic contribution of sFRP2 OE by analyzing cell morphology and cell proliferation rate in ectopic sFRP2 OE and KD condition. OE in WT (sFRP2OE) cells resulted in a dramatic increase in aggregated structures, reminiscent of the initial step in tumorigenesis, and was enhanced in ectopic sFRP2 OE in LFS cells at day 14-17 of OB differentiation (Fig 4A). On the other hand, sFRP2 OE at the MSC stage did not generate aggregated structures (data not shown). OE also increased proliferation during MSC culture and OB differentiation (Figs 4B and Supplementary S4A). sFRP2KD cells had altered cell morphology (not shown) and decreased cell proliferation rate compared to LFS cells (Figs 4C and Supplementary S4B).

**Figure 4.**
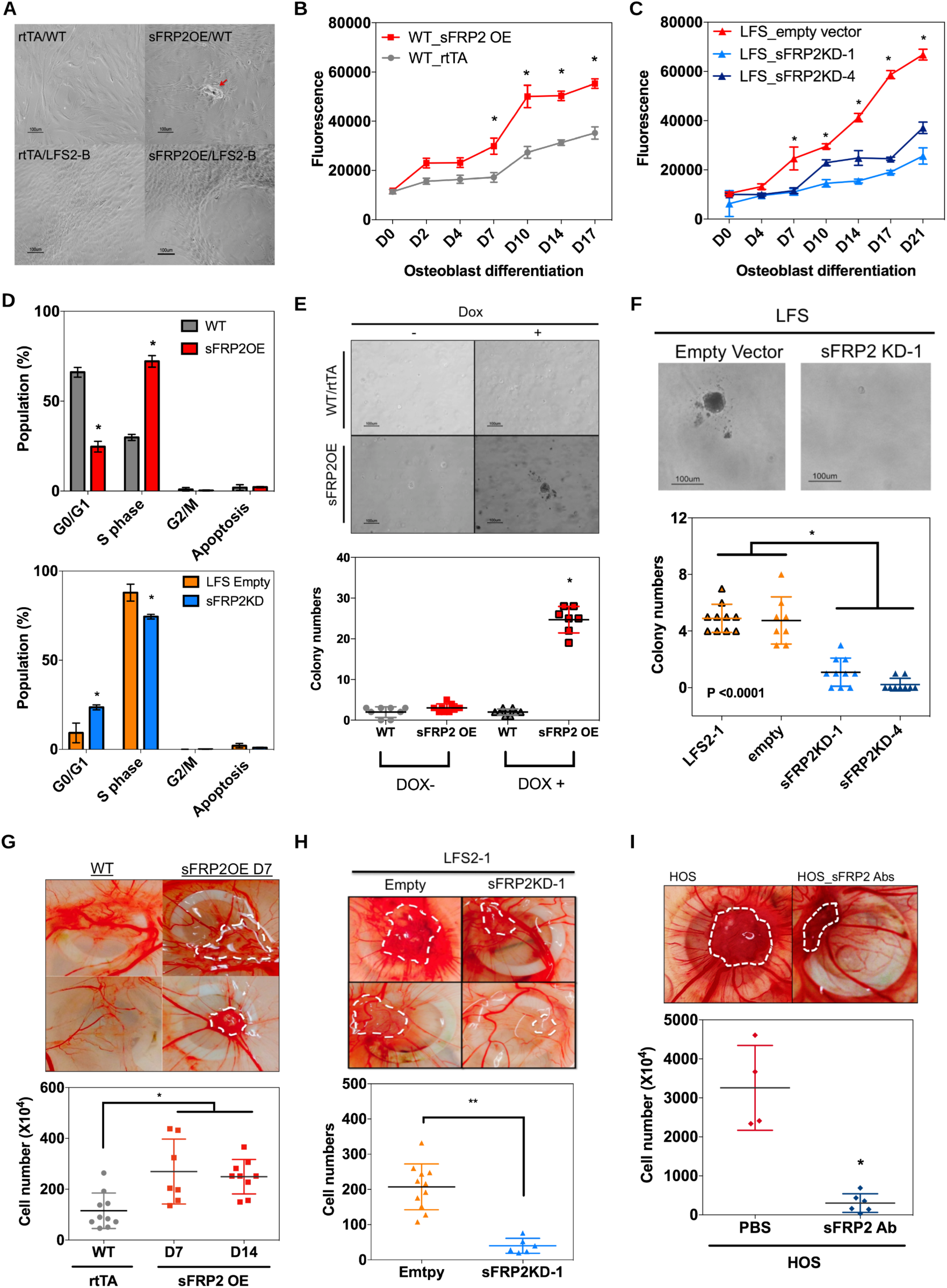
sFRP2 OE in pre-OBs induces cell transformation. **A.** Morphological changes in sFRP2OE cells (Red arrow, aggregated cells). **B.** Cell proliferation analysis (mean±SD, n=4) of sFRP2OE (left). **C.** sFRP2KD (right) using CyQuant Cell assay (fluorescent DNA content; *, p-value<0.0001, Two-way ANOVA). **D.** Cell cycle analysis of WT and sFRP2OE cells using BD BrdU/7-AAD at day 4 of OB differentiation (mean±SD, n=3 independent repeats in duplicate; *, p-value< 0.0001, Two-way ANOVA). **E.** Soft agar colony formation of sFRP2OE and WT cells. **F.** Soft agar colony formation of sFRP2KD and LFS cells. Colonies over 50 µm in size were included in the quantification of colony numbers (bottom panels; *, p-value<0.0001; scale bar, 100 µm). **G.** CAM assay analysis of WT and sFRP2OE cells. **H.** CAM assay analysis of LFS and sFRP2KD cells. **I.** CAM assay of an osteosarcoma cell line (HOS) treated with sFRP2 neutralizing antibody.

It was reported that ectopic sFRP2 expression alters the G2 phase of the cell cycle and suppresses apoptosis (Yamamura et al, 2010). Our analysis of both cell cycle and apoptosis on day 4 of OB differentiation revealed significantly more sFRP2OE cells in Sphase (72.17 ± 3.22 %) than WT cells did (29.82 ± 1.68 %). It implies that sFRP2 OE promotes S phase entry, cell cycle progression and, thus, cell proliferation (Figs 4D and Supplementary S4C). In contrast, KD of sFRP2 resulted in significantly less cell proliferation (Figs 4C and D, and Supplementary S4B and D) with more sFRP2KD cells in G0/G1 (21.51 ± 2.68%) and fewer in S phase (77.36 ± 2.86%) than LFS cells (G0/G1, 8.60 ± 4.32%; S phase, 90.51 ±1.99%)(Figs 4C and D, and Supplementary S4B and D). No statistical difference in apoptotic rates was found among the various groups (Fig 4D). Therefore, sFRP2 OE promotes S phase entry at the pre-OB stage.

Anchorage-independent growth is a hallmark of cancer cell transformation. We used the soft agar colony formation assay to determine the transformation ability of sFRP2. sFRP2OE and LFS cells were able to form oncogenic colonies (Figs 4E and F). Both the number and size of colonies formed by sFRP2OE and LFS cells were significantly greater than that of WT cells. In contrast, sFRP2KD cells formed significantly fewer colonies that LFS cells (Fig 4F). These data clearly demonstrate the tumorigenic ability of sFRP2.

In order to determine the tumorigenicity of sFRP2 OE in LFS and OS cells, we performed *in ovo* CAM assays (Supplementary Fig S4E). This assay provides a convenient and common way to check oncogenicity of sarcoma cells (Fontenot et al, 2013; Sys et al, 2012; Sys et al, 2013). sFRP2OE cells showed increased tumor formation and cell numbers compared to WT cells (Fig 4G). sFRP2KD cells had suppressed tumor formation compared to LFS cells (Fig 4H). Since sFRP2 is a secreted molecule, we wondered if a blocking antibody would reduce sFRP2-mediated tumor formation. Indeed, adding a sFRP2 antibody to HOS cells significantly reduced tumorigenesis (Fig 4I). In serial transplantation analysis, sFRP2OE cells maintained their tumor formation ability (Supplementary Fig 4F) while the antibody in HOS consistently suppressed it over three consecutive transplantations as well (Supplementary Fig 4G). Additionally, using CAM assay, we checked whether FOXM1 inhibition by the small molecule FDI-6 could suppress tumor formation. The result showed that FDI-6 treatment effectively decreased sFRP2 mediated tumor formation (Supplementary Fig 4H). These *in ovo* data demonstrate that sFRP2 OE contributes to LFS mediated tumorigenesis and OS development and that targeting sFRP2 with an antibody suppresses tumor formation.

### Secreted sFRP2 facilitates angiogenesis and tumorigenesis through CYR61

Angiogenesis is a critical component of solid tumor growth. sFRP2 has been shown to function as an angiogenic factor in human triple negative breast cancer, angiosarcoma, and melanoma (Courtwright et al, 2009; Fontenot et al, 2013; Kaur et al, 2016). Whether sFRP2 has any angiogenic properties in OS, however, remains unknown. Because sFRP2 is a secreted protein, we hypothesized that MSCs and pre-OBs in LFS patients could secrete sFRP2 into the bone matrix where it initiates migration and sprouting of endothelial cells, thereby facilitating neoangiogenesis. To investigate this, we conducted an *in vitro* angiogenesis assay; we cultured human vascular endothelial cells (HUVECs) on Matrigel in WT, LFS, or sFRP2OE conditioned medium (CM) for 16 hours and then quantified tube formation rate. As expected, WT CM was not able to significantly induce tube formation (Figs 5A and B). In contrast, LFS and sFRP2OE CM dramatically increased tube formation, as did the addition of recombinant sFRP2 protein to WT CM (Figs 5A and B). Moreover, sFRP2KD CM dramatically decreased tube formation to levels of the negative control and WT CM. This result is consistent with other reports showing the angiogenic function of sFRP2 in angiosarcoma and melanoma (Courtwright et al, 2009; Kaur et al, 2016).

**Figure 5.**
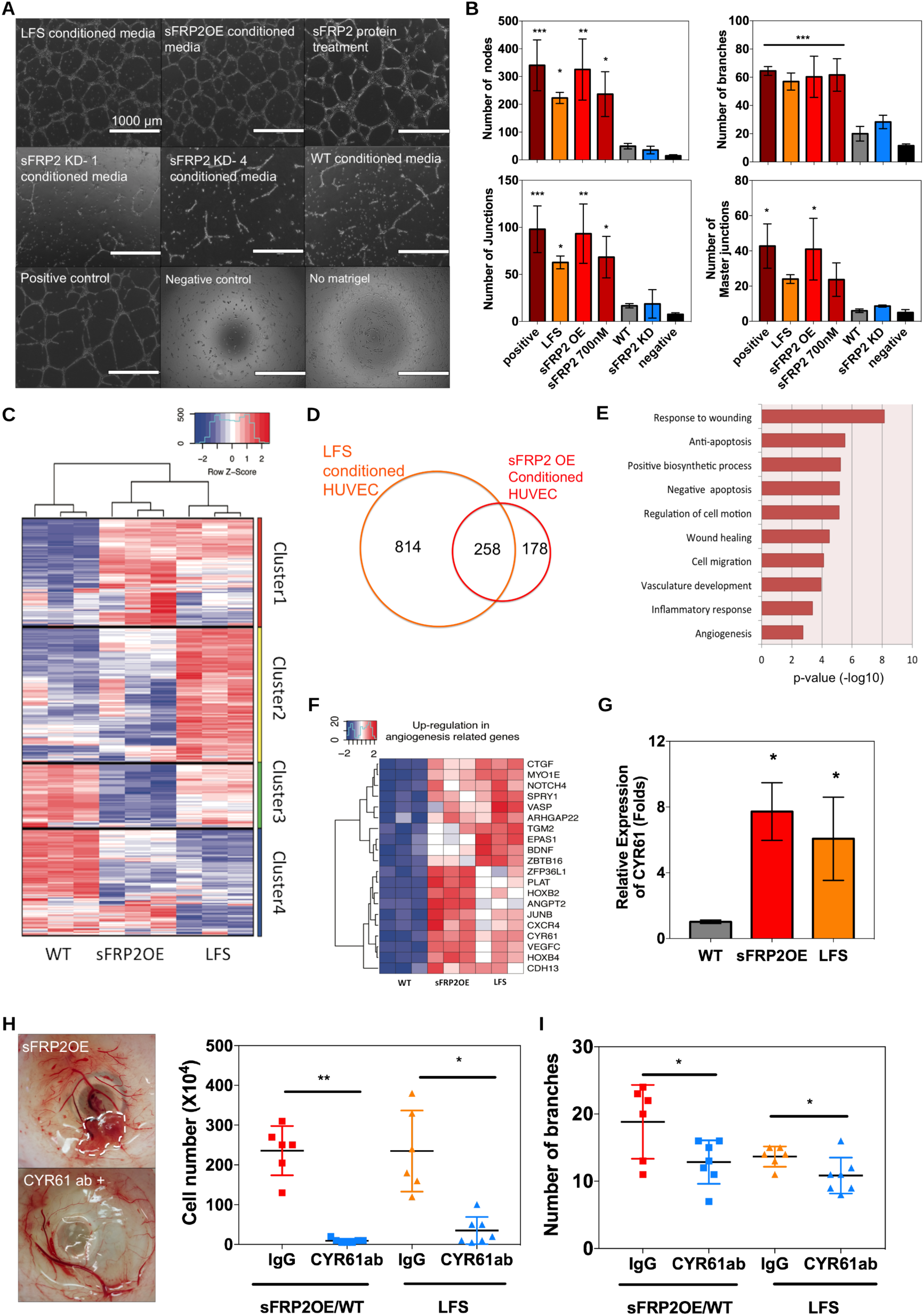
Secreted sFRP2 facilitates angiogenesis. **A.** *In vitro* tube formation of human endothelial cells in CM as indicated. **B.** Quantification (mean±SD, n=3 independent experiments) of tube formation assay (*, p-value<0.05; ** p-value <0.001 ***, p-value<0.0001). **C.** Cluster analysis of the transcriptome of HUVECs in LFS, sFRP2OE and WT CM. **D.** Venn diagram of upregulated DEGs between LFS and sFRP2OE conditions. **E.** DAVID GO functional analysis of upregulated common DEGs in LFS and sFRPOE. **F.** Expression of angiogenic markers in WT, sFRP2OE and LFS CM treated HUVECs. **G.** Q-PCR analysis of CYR 61 expression in endothelial cells after WT, sFRP2OE and LFS CM treatment (mean±SD, n= 2 independent repeats in triplicates; *, p-value< 0.0001). **H.** Tumor formation rate of LFS and sFRP2OE pre-OBs in CAM assay in the presence of CYR61 and control (Rabbit IgG) antibody (*, p-value< 0.005; **, p-value< 0.0001). **F.** Expression of angiogenic markers in WT, sFRP2OE and LFS CM treated HUVECs. **G.** Q-PCR analysis of CYR61 expression in endothelial cells after WT, sFRP2OE and LFS CM treatment (mean±SD, n=2 independent repeats in triplicates; *, p-value<0.0001). **H.** Tumor formation rate of LFS and sFRP2OE pre-OBs in CAM assay in the presence of CYR61 and control (Rabbit IgG) antibody (*, p-value<0.005; **, p-value<0.0001). **I.** Quantification of neovascularization in CAM assay after CYR61 antibody treatment (*, p-value<0.005).

To investigate how secreted sFRP2 increases angiogenesis, we performed RNA-seq profiling of HUVECs after treatment with LFS, sFRP2OE, and WT CM. PCA with triplicate samples from each condition showed that the replicas clustered together but were distinct among the conditions (Supplementary Fig 5A). Unsupervised clustering of the transcriptome profiles showed that sFRP2OE and LFS CM-induced genes clustered more closely to each other than to those of WT CM treated HUVECs (Fig 5C). From unsupervised clustering of the top 500 most varying genes (Variance >0.84), we identified four clusters (1 to 4); Cluster 1 contains genes upregulated in LFS and sFRP2OE compared to WT while Cluster 4 contains genes downregulated in both LFS and sFRP2OE conditions. Cluster 2 and 3 include genes showing different patterns between LFS and sFRP2OE, though LFS and sFRP2OE are closer to each other than to WT based on these 500 genes as indicated by the dendrogram. To identify sFRP2-mediated effects on angiogenesis induction, we focused on Cluster 1 and 4. Using DESeq2 (p.adjusted <0.01), we found 498 common genes (258 upregulated and 240 downregulated) between 1853 LFS and 969 sFRP2OE DEGs (Figs 5D and Supplementary S5B). DAVID functional annotation of the 258 upregulated common genes showed that secreted sFRP2 mainly alters angiogenesis, anti-apoptosis and inflammatory response processes of endothelial cells (Fig 5E). Enrichr GO cellular component analysis of genes linked to angiogenesis-related GO terms were associated with focal adhesion (GO:0005925, p-value=0.000377) (Fig 5F). Interestingly, most upregulated angiogenic genes were either well-known angiogenic markers or pro-cancer genes such as VEGFC, CYR61, VASP, EPAS1, JUNB, and CDH13 (Fig 5F). Recently, it was reported that CYR61, cysteine-rich heparin-binding inducer 61, facilitates angiogenesis and its high expression level is associated with poor prognosis in osteosarcoma (Habel et al, 2015; Liu et al, 2017). We also observed increased expression of CRY61 in endothelial cells treated with sFRP2OE and LFS CM (Figs 5F and G). Thus, we investigated if sFRP2 OE increases angiogenesis and tumor formation through an action of CYR61. We measured the level of neovascularization and tumor size in CAM assay. The results show that CYR61 antibody treatment decreases angiogenesis and tumor formation in the presence of sFRP2 (Figs 5H and I) thus supporting that secreted sFRP2 facilitates angiogenesis of endothelial cells and tumorigenesis, at least in part, through CYR61.

Our RNA-seq analysis revealed the upregulation of anti-apoptosis related genes in HUVECs activated by LFS/sFRP2OE CM (Supplementary Fig S5D). The number of viable HUVECs in sFRP2OE and LFS CM were higher than that in WT CM (Supplementary Fig S5E). These results suggest that secreted sFRP2 facilitates angiogenesis and protects endothelial cells from apoptosis by inducing the expression of angiogenic/oncogenic factors and anti-apoptosis regulators, respectively. Intriguingly, cell cycle-related DEGs in LFS/sFRP2OE CM treated HUVECs were mainly downregulated (Supplementary Figs S5C and F). There was little overlap in cell cycle-related DEGs between LFS/sFRP2OE CM activated HUVECs and in LFS/sFRP2OE OBs (Supplementary Fig S5G). For example, positive cell cycle regulators such as CDC25B, CKS1B, DLGAP5, GTSE1, JAG2, and NUSAP1 were downregulated in LFS/sFRP2OE endothelial cells but were upregulated in LFS/sFRP2OE OBs (Supplementary Fig S5G). Only three genes showed common expression pattern between the two groups; JUNB and MYC, two tumorigenic factors that were upregulated, while KRT18, an intermediate filament gene, was downregulated in both OBs and HUVECs affected by sFRP2 signaling (Supplementary Fig S5G). We speculated that HUVECs in LFS and sFRP2OE CM stopped proliferating and initiated differentiation to generate neo-vasculature. This result emphasizes that the function of sFRP2 is context dependent.

### Targeting sFRP2 OE effectively suppresses OS development *in vivo*

To examine if suppression of sFRP2 expression in LFS has any therapeutic effect on OS development *in vivo*, we subcutaneously injected LFS and sFRP2KD cells at day 7 of OB differentiation into Nu/Nu mice. Tumors rapidly formed in mice injected with LFS3-A (at 2 weeks post-injection) or LFS2-B cells (at 3 weeks post-injection) (Supplementary Fig S6A). We attribute the difference in tumor formation timing to lower sFRP2 expression in LFSB-2 than in LFS3-A (Fig 1B). In contrast, mice injected with sFRP2KD cells either did not form tumors or tumors were of very small size (Figs 6A and Supplementary S6A). Histological analysis revealed that LFS tumors were composed of large, poorly differentiated malignant tumor cells with mitotic features and large areas of necrosis reminiscent of tissue morphology seen in human osteoblastic or fibroblastic OS (Figs 6B and Supplementary S6B) (Daft et al, 2013). A few of the small tumors in the sFRP2KD group showed some neoplastic features, while others showed induction of fibrous tissues (Figs 6B and Supplementary S6B). Interestingly, LFS tumors showed lower levels of AP, a marker of mature OBs, than tumors formed from sFRP2KD cells (Fig 6B). Despite higher β-catenin expression, β-catenin was localized to the cytosol, granules and cell membrane in the sFRP2KD group (Fig 6B). This finding augments the idea that sFRP2-mediated tumorigenesis is not directly linked to hyperactive WNT signaling. Furthermore, we found more vasoformation as assessed by CD31 endothelial marker expression in LFS cell-induced tumors than in the sFRP2KD group (Figs 6C and D). These histological analyses support the idea that sFRP2 OE facilitates neovascularization and tumorigenesis. Similarly to the sFRP2KD group, targeting sFRP2 in malignant HOS/MNNG OS cells reduced OS tumor growth Supplementary Fig S6C). Additionally, we checked the expression of signature genes altered by sFRP2 OE by RNA-seq analysis in these primary tumors (Figs 6E and Supplementary S6D). Oncogenic CYR61 and FOXM1 were only differentially expressed in primary tumors between sFRP2KD and empty vector groups of OS Tumor suppressor genes while CADM1 and CADM4 were significantly upregulated in sFRP2KD OBs and sFRP2KD primary tumors compared to parental LFS and OS. (Figs 6E and Supplementary S6D).

**Figure 6.**
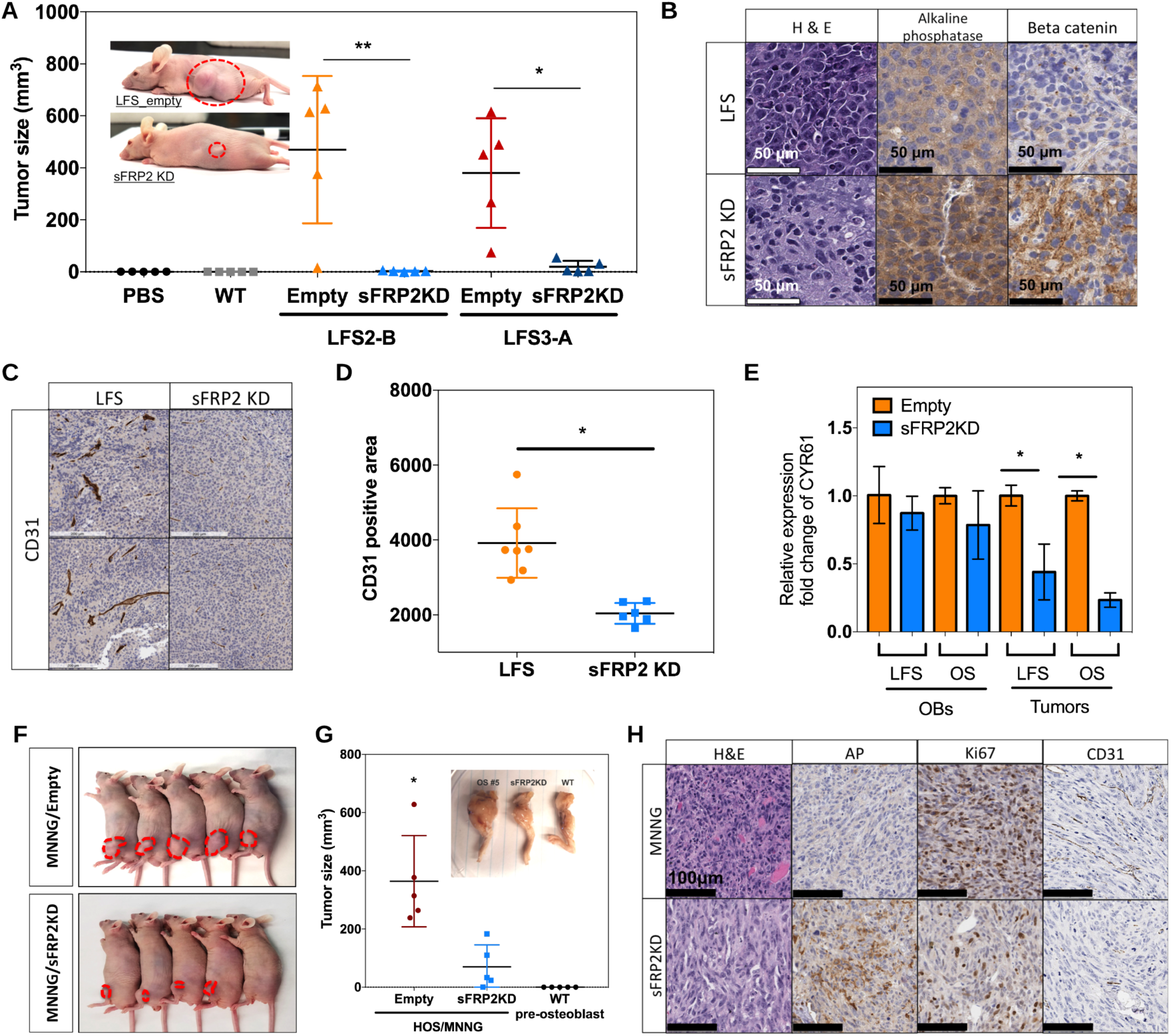
Targeting sFRP2 in LFS pre-OBs and OS cell line suppresses osteosarcoma development *in vivo*. **A.** Tumor formation of LFS and sFRP2KD cells in Nu/Nu mice four weeks after cell injection (tumor volume = 3.14/6*L*W*H). Each dot, square or triangle represents an individual analyzed mouse. (n=5; *, p-value<0.001, **, p-value< 0.0001. **B.** Immunohistochemistry analysis of LFS and sFRP2KD cell-induced tumors. **C.** CD 31 positive area in LFS and sFRP2KD cell-induced tumors. **D.** Quantification of CD31 expression in tumors. Each dot or square represents an individual analyzed induced tumor/mouse (*, p-value<0.05). **E.** Expression of CYR61 altered by sFRP2 OE in LFS OBs and OS cells and subcutaneous primary tumors of LFS and OS cells (n=4; *, p-value <0.05). **F.** Intratibial tumor formation of osteosarcoma (MNNG) and sFRP2KD groups. **G.** Tumor size 4 weeks after injection. Each dot or square represents an individual analyzed induced tumor/mouse (*, p-value<0.001). **H.** Representative immunohistochemistry analysis of OS and sFRP2KD tumors.

We next performed intratibial injection in mice (Husmann et al, 2013; Vormoor et al, 2014; Yuan et al, 2009) to investigate the orthotopic effect of sFRP2 depletion in OS development. First, depletion of sFRP2 in the malignant OS cell line, HOS/MNNG, significantly reduced tumor incidence and size (Figs 6F and G). Primary lesions showed characteristics of primary human OS with osteolytic, mitotically active undifferentiated cells, as well as wide bone destruction (Figs 6H and Supplementary S6E). Microscopically, HOS/MNNG tumors showed less AP activity, more proliferating cells (Ki67^+^) and more neovascularized areas (CD31^+^) than the sFRP2-depleted HOS/MNNG tumors (Fig 6H). The bone structure in the sFRP2KD group was relatively intact although the tumors still contained undifferentiated tumorigenic cells in the proximal tibia (Fig 6G and Supplementary S6E). We also performed intratibial injections with LFS and sFRP2KD cells. In the LFS group, two mice generated tumors in the tibia five months post-injection, and one mouse formed a tumor on the flank of the body but not intratibially (Fig 6F). This might have been caused by a failure of proper intratibial injection. Three mice died within three months of injection with no visible exterior tumor formation but with clear signs of weight loss. The actual cause of death was undetermined. In stark contrast, the sFRP2KD group did not form any osteosarcomatous lesions (Fig EV6F). Conclusively, the *in vivo* xenograft study unequivocally demonstrates that depletion of sFRP2 in LFS mediated tumorigenesis and OS development has potential therapeutic benefits.

## Discussion

Here we show that LFS patient-specific iPSCs and their MSCs and OBs provide a novel platform to understand the early development of OS and identify and evaluate a potential therapeutic target, sFRP2, for this devastating cancer. Using molecular and functional experiments, traditional *in vivo* studies, and bioinformatic analyses we found that sFRP2 expression significantly contributes to LFS-mediated tumorigenesis and OS development by facilitating angiogenesis and cell proliferation. Most significantly, we demonstrated that targeting sFRP2 in LFS and OS has a potential therapeutic application.

### sFRP2 suppressed canonical WNT pathway is associated with abnormal OB differentiation, but not cell transformation

We found that sFRP2 is upregulated in LFS patient cells and OS cell lines harboring *TP53* mutations. We also showed that sFRP2 exerts oncogenicity in MSCs/OBs both *in vitro* and *in vivo*. These findings conflict with the hyper-WNT signaling paradigm in cancer biology. Generally, sFRP2 is widely known as an antagonist of the WNT pathway due to its ability to sequester WNT ligand from the frizzled receptor and is predominantly considered to be a tumor suppressor. Indeed, promoter hypermethylation of sFRP2 is frequently found with increased WNT signaling in many cancers such as colorectal, bladder, and gastric cancer and carcinoma (Marsit et al, 2005; Nojima et al, 2007; Silva et al, 2014). However, in an LFS context, sFRP2 functions as an oncogenic factor. We consistently observed that sFRP2 OE suppressed WNT signaling and that the majority of β-catenin remained localized to the cytoplasm. Our data also indicate that lower β-catenin expression is associated with initiation of abnormal OB differentiation and is maintained to the terminal stage of osteosarcomagenesis (*in vivo* study). This finding is consistent with a report of inactive β-catenin and WNT signaling in high-grade human OS (Cai et al, 2010). This study adds an evidence of the complexity of WNT pathway regulation and its diverse outcomes in different conditions.

The mechanism by which sFRP2 exerts its role is unknown in OS development and remains elusive, context-dependent and even controversial in many other cancers. A recent paper described an age-related increase in sFPR2 expression in fibroblasts that activated a multi-step signaling cascade in melanoma cells resulting in a reduction of β-catenin and microphthalmia-associated transcription factor (MITF) expression, and consequently the loss of APE1. The absence of APE1 affected DNA damage response of the cells, increased metastasis and rendered the cells more resistant to chemotherapy (Kaur et al, 2016). In our study, we observed that sFRP2 OE suppresses β-catenin, but MITF or APE1 expressions were not altered. Instead, we found that sFRP2OE in OBs upregulates FOXM1 in OBs as an autocrine factor and CRY61 in endothelial cells as a paracrine factor in the β-catenin independent way. Therefore, we concluded that sFRP2 associated osteosarcomagenesis develops through a mechanism different than that in melanoma.

### sFRP2 OE in LFS contributes to an intrinsic oncogenic molecular profile

Oncogenesis is commonly attributed to abnormal differentiation, aberrant cell adhesion, dysregulated cell cycle and overproliferation. Our transcriptomic analysis of sFRP2OE and sFRP2KD cells undoubtedly showed that sFRP2 plays multiple functions during abnormal differentiation and tumorigenesis in LFS OS development. Specifically, sFRP2 OE changes the cell cycle profile and increases expression of cell proliferation-related genes such as AURKB and FOXM1. AURKB regulates chromosomal segregation during mitosis and meiosis, and aberrant expression of AURKB is observed in several cancers (Gonzalez-Loyola et al, 2015; Smith et al, 2005). FOXM1 is also found overexpressed in many cancers (Huang et al, 2014; Radhakrishnan et al, 2006). Using our LFS OB model, in *in ovo* CAM assay, we found that inhibition of FOXM1 impedes OS development (Fig. S4 H). FOXM1 regulates transcription of cell cycle and proliferation genes (Laoukili et al, 2005; Wang et al, 2008), including subunits of the SCF ubiquitin ligase complex, Skp2 and Cks1. These two genes ubiquitinate P21 (Cip1) and P27 (Kip1), regulators of cell cycle progression into S phase, for degradation (Petrovic et al, 2008). Therefore, the observed increase in cell proliferation and numbers of cells in S phase of LFS and sFRP2OE cells may be explained by sFRP2 OE induced FOXM1. One may assume that sFRP2 depletion in LFS (sFRP2KD) cells should reduce FOXM1 expression. However, this is not what we observed. We explain this by the fact that FOXM1 autoregulates its expression (Cheng et al, 2014; Halasi & Gartel, 2009). Thus, once FOXM1 expression is induced, its expression is maintained and sFRP2 depletion has no apparent effect. Nevertheless, FOXM1 inhibition by way of FDI-6 small molecule treatment effectively decreased tumor formation (Supplementary Fig S4H). It is noteworthy that FOXM1 upregulation was correlated with poor prognosis of OS patients (Fan et al, 2015). This fits well with our own observation that high sFRP2 expression in OS tissues is linked to decreased patient survival (Figs 1F and Supplementary S1E).

Additionally, we found that commonly downregulated genes between sFRP2OE and sFRP2KD cells include cell adhesion genes such as CADM1 and CADM4. They are known tumor suppressors in various malignancies, such as brain, prostate, breast, and pancreatic cancer (Nagata et al, 2012; Nowacki et al, 2008; Sakurai-Yageta et al, 2009; Vallath et al, 2016). The significance and role of CAMD1 and CADM4 downregulation in OS development is unknown. A possibility is that their downregulation is linked to anchorage-independent cell growth and cell egress from the bone environment resulting in metastasis. This warrants future investigation. Taken together, abnormal expression of these specific genes (AURKB, FOXM1, CADM1, and CADM4) clearly indicates the oncogenic property of sFRP2 overexpression in OS.

### Secreted sFRP2 promotes angiogenesis and tumorigenesis associated with upregulation of CYR61 in endothelial cells

Angiogenesis is a critical hallmark of cancer progression. In this study, we showed that secreted sFRP2 facilitates angiogenesis *in vitro* and *in vivo*. Global transcriptome analysis revealed up-regulation of angiogenesis-related and anti-apoptotic genes in endothelial cells grown in LFS, sFRP2 cell-conditioned and sFRP2-supplemented media. Angiogenesis-related genes such as CDH13, EPAS1, and CYR61 are highly associated with focal adhesion and tumorigenesis. Particularly, CYR61, a secreted extracellular matrix protein, is a pro-angiogenic/tumorigenic molecule validated in many cancers (Chen et al, 2016; Leask & Abraham, 2006). Interestingly, CYR61 has been found to mediate specific functions in different types of cells through binding to distinct integrins during angiogenesis (Leu et al, 2002; Park et al, 2015) and metastasis (Sun et al, 2008). CYR61 has been reported to be overexpressed in OS compared to normal bone tissues and that its depletion causes inhibition of neoangiogenesis in the developing tumor (Habel et al., 2015). We found that CYR61 inhibition prevents sFRP2 mediated OS development in *in ovo* CAM assay (Figs 5H and I). Collectively, the sFRP2-CYR61 axis in LFS may alter the extracellular matrix of endothelial cells thus promoting angiogenesis, tumorigenesis and possibly metastasis. Surprisingly, secreted sFRP2 did not increase expression of cell-cycle related genes in endothelial cells. This is in contrast to LFS and sFRP2OE OBs.

Overall, our data provide new insights into the role of sFRP2 in OS initiation and development. Using the powerful tool of iPSC-based modeling, we show that sFRP2 has a dual function. sFRP2 increases the proliferative capacity of OS cells, and, can also, promote angiogenesis through cell adhesion/cell matrix factors; sFRP2-FOXM1 in LFS OB/OS and sFRP2-CYR61 in endothelial cells contribute to OS development. We did not investigate the effect of sFRP2 in metastasis of LFS/OS, however, a recent report showed that sFRP2 potentially plays a crucial role in metastatic OS, where its expression becomes even more enhanced in low metastatic mouse OS cells and contributes to invasiveness and metastasis without affecting metastatic OS cell proliferation (Techavichit et al, 2016). Detailed future exploration of the sFRP2-FOXM1 and sFRP2-CYR61 axes during OS development and metastasis will provide additional mechanistic insight and help develop novel therapeutic strategies to treat LFS and OS.

## Materials and Methods

### WT and LFS iPSCs and *in vitro* differentiation into OBs

WT and LFS iPSCs were established in Dr. Ihor Lemischka’s laboratory as previously reported (Lee et al., 2015). Three LFS patients and two WT fibroblasts were reprogrammed using integration-free Sendai virus for delivery of the reprogramming factors. Established iPSC lines were cultured in hESC medium consisting of DMEM/F12 supplemented with 20% Knockout serum replacement, 10ng/ml basic FGF (Invitrogen) on mouse feeder cells (CF1) (Invitrogen) or in mTeSR (STEMCELL technology) on Matrigel coated plates. The iPSCs were differentiated to MSCs using well-established MSC differentiation medium (DMEM supplemented with 10% Knockout serum replacement, 5 ng/ml FGF2 and 5 ng/ml PDGF-AB [PeproTech]). After 3 weeks of differentiation, cells were sorted for CD105^+^ and CD24^−^ cells using a BD Aria ll at the Mount Sinai Flow Cytometry Shared Facility. Sorted cells were expanded in MSC medium (DMEM supplemented with 10% FBS). WT and LFS iPSC-derived MSCs were differentiated in osteoblast differentiation medium (α-MEM supplemented with 10% FBS, 0.1 mM dexamethasone, 10 mM β-glycerol phosphate, and 200 mM ascorbic acid) at 37 °C in a 95% air, 5% CO_2_ humidified incubator (Bilousova et al, 2011). The information on differentiated cell lines utilized in this study is in Supplementary Table S1.

### Osteosarcoma cell lines

Human osteosarcoma cell lines were obtained from Dr. Robert Maki (Icahn School of Medicine) and Dr. Nino Rainusso (Texas Children’s Hospital, Houston, TX 77030-2399). HOS, HOS/MNNG, HOS/143B cells were culture in DMEM supplemented with 10% FBS. SJSA-1 and USO2 cells were cultured in RPMI supplemented with 10% FBS at 37°C in a 95% air, 5% CO_2_ humidified incubator.

### Quantitative real time PCR

Total RNA (1mg) was isolated with Trizol Reagent (Life Technologies). cDNAs was generated using the SuperScript^®^ III Reverse Transcriptase kit (Life Technologies). SYBR Green Real-Time PCR Master Mixes kit (Life Technologies) was used for real-time quantitative PCR with a Roche Applied Science LightCycler 480 Realtime PCR machine (Applied Biosystems) and the primer sets listed in Supplementary Table S4.

### Western blotting

Total cellular protein was extracted using Radio-Immunoprecipitation Assay buffer (RIPA) and boiled to denature all proteins. Cytosolic and nuclear proteins were isolated using Nuclear/Cytosol Fractionation kit (Biovision.Inc). Cell extracts were subjected to sodium dodecyl sulfate–polyacrylamide gel electrophoresis (SDS-PAGE; 10% polyacrylamide gels); and then transferred to polyvinylidene difluoride membranes (Bio-Rad) and blocked with 5% nonfat milk (Labscientific) in TBS–Tween 20 (TBST). Membranes were incubated with primary antibodies overnight at 4 °C. After 3 washes with TBST, membranes were probed with a horseradish peroxidase–conjugated secondary antibody in either 5% nonfat milk or 5% BSA/PBS solution for 1 hr at room temperature. Proteins were detected using SuperSignal West Pico Chemiluminescent Substrate (Thermo).

### CRISPR/Cas9

We designed CRIPSR guide RNAs to target the intronic region adjacent to P53 (G245D) using the web-based tool of the Zhang Lab at MIT (http://crispr.mit.edu:8079). The guide RNA sequences were ligated into the pX335 CRISPR plasmid (Cas9 nickase; Addgene) (Supplementary Table S5). WT iPSCs-derived MSCs were seeded and transfected with 60 µg donor vector (containing G245D) and 5 µg CRIPSR plasmids using Lipofectamine 3000 (Thermo Fisher Scientific). In order to check for proper genome modification, genomic DNA was isolated from targeted clones, the region spanning the target site amplified by PCR using the primers targeting P53 (G245D). The amplified fragment was run on an agarose gel, gel extracted and Sanger sequenced.

### AP and Alizarin Red S staining

AP and Alizarin Red S staining were examined using the alkaline phosphatase staining kit II (00-0055, Stemgent) and Alizarin Red S indicator (Ricca Chemical Company), respectively, following the manufacturers’ recommendations.

### Luciferase reporter assay

BMP reporter (Addgene, plasmid #16495) or WNT Plasmids (Addgene: plasmids #12456 and #12457) were co-transfected with Renilla Luciferase Control reporter (pRLSV4) into hiPSC-derived MSCs, and cells were grown in osteoblast differentiation medium for 14 days. Cells were washed with PBS, lysed in Passive Lysis Buffer and reporter activity measured according to the manufacturer’s instruction (Dual-luciferase reporter assay kit, Promega). Luciferase activities were normalized to the intensity of the Renilla Luciferase Control reporter. Relative activities were calculated in relation to the intensity of no treatment groups.

### β-Catenin immunostaining and quantification

Day 4 pre-osteoblast cells were washed with PBS, fixed with 4% formaldehyde/PBS for 15 minutes, washed again with PBS and blocked with 5% BSA/PBS for 60 mins. The blocking solution was aspirated and applied with diluted conjugated antibody (Rabbit Alexa594 anti-beta catenin, Abcam) overnight at 4°C. Nuclei were stained with DAPI (4′, 6-diamidino-2-phenylindole). Images using the same settings were taken with a fluorescence microscope (Ix51 inverted Microscopy, Olympus). The intensity of β-catenin presented in the figures was quantified using Image J and normalized to the number of nuclei present in the field of view.

### CyQUANT NF fluorescence proliferation assay

Equal numbers of MSCs were plated and induced to differentiate to osteoblast. Cells were harvested at different time points to obtain OBs representing various differentiation stages (D0 to D17). Cells were washed with HBSS buffer and dispensed in 100ul of 1x dye binding solution into wells of a microplate with the cells and were incubated at 37°C for 30 minutes according to the manufacturer’s instructions (Invitrogen). The fluorescence intensity of each sample was detected and measured using a fluorescence microplate reader with excitation at ~485nm (Model 2300, PerkinElmer).

### Cell cycle analysis

Osteoblasts at day 2 of differentiation were labeled with BrdU (1mM solution) for 24 hours. BrdU-pulsed cells were collected, fixed with BD cytofix/cytoperm buffer and incubated with FITC-BrdU antibody and 7-ADD solution according to the manufacturer’s instruction (BD Biosciences). Cell cycle and DNA synthesis activities were determined by analyzing the correlated expression of total DNA and incorporated BrdU levels using flow cytometry (BD LSR II, BD Biosciences).

### CAM xenograft assay

CAM assay was conducted as described previously (Sys et al., 2013). Briefly, 1×10^6^ differentiated osteoblasts (D7 or D14) were mixed with Matrigel and implanted onto the CAM of 8-day-old chick embryos (Charles Rivers Laboratories). After 4 days, tumors were resected, minced, dissociated into a single cell suspension with type 1A collagenase (Roche) and plated on dishes. The total number of tumor cells was counted using a hemocytometer.

### Angiogenesis/tube formation assay

*In vitro* Angiogenesis Assay kit (Abcam) was used for the tube formation assay. Briefly, we added 50 µl of thawed extracellular matrix solution (Matrigel) to each well of a pre-chilled 96-well plate. The plate was incubated for 1 hour at 37°C. Thereafter, HUVECs (passage 6 to 8) were harvested and plated at 2x 10^4^ cells per well. HUVECs were cultured in conditioned medium for 16 hours at 37 °C, 5% CO_2_ and tube formation was captured by taking images with an EVOS FL Cell Imaging system (ThermoFisher). The CM was collected from Day 7 *in vitro* OB cultures of WT, sFRP2OE, and LFS cells. Images were analyzed with Image J Angiogenesis Analyzer tool. This tool quantifies the tube formation images by extracting characteristic information of the formed tube meshes and branches (Chevalier et al, 2014).

### Mouse study

Male 8-week-old nude/nude mice were ordered from Charles River and used for intratibial and subcutaneous injections of LFS and OS cells. Since general osteosarcoma frequency is higher in the male than the female population and this study did not focus on investigating variations, including of tumor incidence, between males and females, we only included male mice. The sample size was determined based on power calculation (GPOWER software; with one-way ANOVA) and our estimation on surgical failure (Dell et al, 2002). Mice were housed in the animal facility of the Icahn Medical Institute building at the Icahn School of Medicine. Mice were given ad libitum access to water and standard rodent chow in a temperature-controlled environment. All animal use was reviewed and approved by the Institutional Animal Care and Use Committee (IACUC) at the Icahn School of Medicine at Mount Sinai. All experiments were conducted according to federal, state, and local regulations under the supervision of four full time trained veterinarians of the Center for Comparative Medicine and Surgery at the Icahn School of Medicine at Mount Sinai.

Mouse xenograft tumor specimens were prepared and analyzed by pathologists at Histowiz (https://www.histowiz.com). Tumors were fixed in 1:10 dilution-buffered Formalin; dehydrated and embedded in paraffin; and sectioned and processed for hematoxylin and eosin (H&E) staining, Picro Sirus Red staining (STPSRPT, American Master Tech) and Von Kossa Staining (KTVKO, American Master Tech). Samples were processed, embedded in paraffin, and sectioned at 4 µm. IHC was performed on a Bond Rx autostainer (Leica Biosystems) with heat mediated antigen retrieval using standard protocols. The sections were deparaffinized and incubated with the primary antibody CD31 (PA0250, Leica Biosystems), Ki-67 (PA0230, Leica Biosystems), Alkaline phosphatase (111871AP, Proteintech), beta catenin (8480, Cell Signaling Technologies) and sFRP2 (PA529590, Thermo Fisher) antibodies. The sections were lightly counterstained using hematoxylin. All slides were scanned with Aperio AT2 (Leica Biosystems) at 40x magnification.

### Human OS tissue microarray

The OS tissue microarray set was obtained from the Pathology Department (Dr. Elizabeth Demicco). It was originally generated for a retrospective study (identification of novel biomarkers in sarcomas,) using Mount Sinai Medical Center-archived formalin-fixed paraffin embedded tissues, which are no longer needed for patient sarcoma diagnosis. The use of the OS microarray set and data analysis was approved by the Icahn School of Medicine at Mount Sinai’s Program for the Protection of Human Subjects and the Institutional Review Board. Immunohistochemistry (IHC) for OS tissue microarray analysis was performed by pathologists at Histowiz (https://www.histowiz.com). Use of sFRP2 antibody for IHC was optimized using mouse muscle (negative control) and human prostate tissue (positive control). Subsequently, the antibody at the optimal dilution of 1:75 was used on the tissue microarray comprising 151 patient OS samples.

### RNA-Seq data analysis

cDNA library preparation and sequencing were performed at Novogen (http://www.novogen.com/home). Briefly, RNA quality was determined with an Agilent Bioanalyzer (RNA integrity number [RIN] > 7.0 for all samples). mRNA was purified from total RNA using poly-T oligo-attached magnetic beads, and double-stranded cDNA synthesized using random hexamers, M-MuLV Reverse Transcriptase (RNase H), and DNA Polymerase I. NEBNext Adaptor with hairpin loop structure were ligated to the cDNA in preparation for hybridization. In order to select cDNA fragments of preferentially 150~200 bp in length, the library fragments are purified with AMPure XP system (Beckman Coulter, Beverly, USA). Library of 150-200 bp fragments at an effective concentration of > 2nM was loaded onto an Illumina HiSeq 4000 for sequencing.

Sequence reads were aligned to human transcript reference (RefSeq Genes, hg19) for the expression analysis at gene levels using TopHat (Trapnell et al, 2009) and HTSeq (Anders et al, 2015). The raw counts of reads aligned to each gene were further normalized by Reads Per Kilobase of transcript per Million (RPKM) as the gene expression abundance measurement. For the original dataset, we used linear regression to identify differentially expressed genes independent of time (D0, D7, D14, and D17) among the different conditions (WT, sFRP2OE, LFS, or sFRP2KD) *G* ~ *time* + *condition*. The p-value cutoff (p<0.001) was validated by permutation test (FDR<0.01). For the HUVEC dataset, DESeq2 was used to compare gene expression among WT, LFS, and sFRP2OE and adjusted p-value cutoff (p<0.01) was applied to define differentially expressed genes.

### Statistical analysis

Student’s t-test (two-sided) was applied, and changes were considered statistically significant for p<0.05. For the comparison of more than two groups, a one-way or two-way (if data include different time points) ANOVA was applied. Survival in human subject and mouse experiments was represented with Kaplan-Meier curves, and significance was estimated with the log-rank test (Prism GraphPad). The data shown in the bar and dot graphs are the mean ±SD of at least three biological replicates. Statistical analysis was conducted using the Microsoft Excel or GraphPad Prism software packages.

## Accession numbers

The RNA-seq data are deposited at GEO under assigned accession numbers GSE102729 and GSE102732. The histological data is available on Histowiz.com, and direct links will be provided upon proper requests and use agreements.

Resources and reagents will be provided upon proper requests and use agreements. According to regulations we do not share patient or clinical information linked to the OS Tissue Microarray set used beyond what is provided in this paper (See Supplementary Table S2).

## Supplementary Material

Supplementary Fig S1 shows association of sFRP2 overexpression with OS development.

Supplementary Fig S2. shows global gene expression profiles of sFRP2OE and sFRP2KD cells.

Supplementary Fig S3 shows suppression of OB differentiation by sFRP2 overexpression.

Supplementary Fig S4 shows initiation of cell transformation by sFRP2 overxpression.

Supplementary Fig S5 show RNA-seq data of sFRP2 activated endothelial cells.

Supplementary Fig s6 shows reduced ostesarcomagenesis upon targeting sFRP2 in LFS and OS cell lines.

Supplementary Table S1 lists the characterizations of WT and LFS cells.

Supplementary Table S2 lists the patient data linked to the OS tissue microarray.

Supplementary Table S3 lists overlaps of DEGs.

Supplementary Table S4 list the PCR and qPCR primers.

Supplementary Table S5 lists recombinant DNAs, antibodies and recombinant proteins.

## Acknowledgements

We dedicate this work to the memory of Ihor R. Lemischka, a scientist, visionary, pioneer, and mentor, who is internationally recognized for his contributions to stem cell research working at the intersection of genomics, systems-level analyses, and cellular and molecular mechanisms of cell fates and disease states. We thank Drs. Robert Maki and Nino Rainusso for providing OS cell lines, Dr. Marion D. Zwaka for sharing shRNAs to P53, and Drs. Zichen Wang and Avi Ma’ayan with help in RNA-sequencing analysis of preliminary data, and Dr. Avi Ma’ayan for reading and input on the manuscript.

This work was supported by a NCI Pre- to Postdoctoral Transition Award (1F99CA212489) and funds provided by the Graduate School of Biomedical Sciences at the Icahn School of Medicine to H.S.K, a NIH Pathway to Independence Award (R00CA181496) and a Cancer Prevention Research Institute of Texas (CPRIT) Award RR160019 to D.F.L., NIH grant (5R01GM078465) to I.R.L, and the Empire State Stem Cell Fund through the New York State Department of Health (NYSTEM) to I.R.L. (C024410).

Author contributions: H.S.K designed, carried out experiments, interpreted data and wrote the manuscript. S.Y. and J.Z. aided in the bioinformatics analysis, Y.X.Y., J.M.B., A.M.G., M.G.D. and J.S. helped perform experiments and provided advice. E.G.D. provided OS tissue microarrays, performed histological analysis and interpreted data. K.A.M. provided advice, reagents and aided in mouse xenograft study. I.R.L., D.F.L and C.S guided and supervised the project and wrote the manuscript. All authors provided input and helped edit the manuscript.

The authors declare no competing financial conflicts of interest.

